## Supplemental Information for "Bayesian inference of relative fitness on high-throughput pooled competition assays"

### Supplementary Materials

#### Primer on Variational Inference

In this section, we will briefly introduce the idea behind variational inference. Recall that any Bayesian inference problem deals with the joint distribution between observations $\underline{x}$ and unobserved latent variables $\underline{\theta}$. This joint distribution can be written as the product of a distribution of the observations $\underline{x}$ conditioned on the $\underline{\theta}$ and the marginal distribution of these latent variables, i.e.,

$$\pi(\underline{x}, \underline{\theta}) = \pi(\underline{x} \mid \underline{\theta})\pi(\underline{\theta}). \tag{S1}$$

A Bayesian inference pipeline's objective is to compute the latent variables' posterior probability given a set of observations. This computation is equivalent to updating our prior beliefs about the set of values that the latent variables take after taking in new data. We write this as Bayes theorem

$$\pi(\underline{\theta} \mid \underline{x}) = \frac{\pi(\underline{x} \mid \underline{\theta})\pi(\underline{\theta})}{\pi(\underline{x})}. \tag{S2}$$

The main technical challenge for working with Equation S2 comes from the computation of the denominator, also known as the *evidence* or the *marginalized likelihood*. The reason computing this term is challenging is because it involves a (potentially) high-dimensional integral of the form

$$\pi(\underline{x}) = \int \cdots \int d^K\underline{\theta}\ \pi(\underline{x}, \underline{\theta}) = \int \cdots \int d^K\underline{\theta}\ \pi(\underline{x} \mid \underline{\theta})\pi(\underline{\theta}), \tag{S3}$$

where $K$ is the dimesionality of the $\underline{\theta}$ vector. Here, the integrals are taken over the support—the set of values valid for the distribution—of $\pi(\underline{\theta})$. However, only a few selected distributions have a closed analytical form; thus, in most cases Equation S3 must be solved numerically.

Integration in high-dimensional spaces can be computationally extremely challenging. For a naive numerical quadrature procedure, integrating over a grid of values for each dimension of $\underline{\theta}$ comes with an exponential explosion of the number of required grid point evaluations, most of which do not contribute significantly to the integration. To gain visual intuition about this challenge, imagine integrating the function depicted in Figure S1. If the location of the high-density region (dark peak) is unknown, numerical quadrature requires many grid points to ensure we capture this peak. However, most of the numerical evaluations of the function on the grid points do not contribute significantly to the integral. Therefore, our computational resources are wasted on insignificant evaluations. This only gets worse as the number of dimensions increases since the number of grid point evaluation scales exponentially.

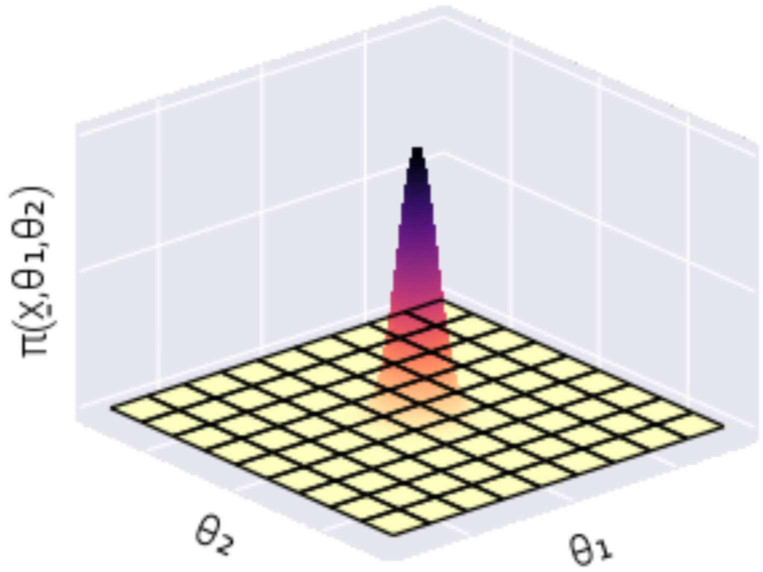

**Figure S1. High-dimensional numerical quadrature does not scale with dimensionality.** Schematic depiction of the problem with naive numerical quadrature to integrate over an unknown density. While the density is concentrated on the dark peak, most of the evaluations over the $x_1 - x_2$ grid do not contribute to the value of the integral

Modern Markov Chain Monte Carlo algorithms, such as Hamiltonian Monte Carlo, can efficiently perform this high-dimensional integration by utilizing gradient information from the target density Betancourt [6]. Nevertheless, these sampling-based methods become prohibitively slow for the number of dimensions our present inference problem presents. Thus, there is a need to find scalable methods for the inference problem in Equation S2.

Variational inference circumvents these technical challenges by proposing an approximate solution to the problem. Instead of working with the posterior distribution in its full glory $\pi(\underline{\theta} \mid \underline{x})$, let us propose an approximate posterior distribution $q_\phi$ that belongs to a distribution family fully parametrized by $\phi$. For example, let us say that the distribution $q_\phi$ belongs to the family of multivariate Normal distributions such that $\phi = (\underline{\mu}, \underline{\underline{\Sigma}})$, where $\underline{\mu}$ is the vector of means and $\underline{\underline{\Sigma}}$ is the covariance matrix. If we replace $\pi$ by $q_\phi$, we want $q_\phi$ to resemble the original posterior as much as possible. Mathematically, this can be expressed as minimizing a "*distance metric*"—the Kullback-Leibler (KL) divergence, for example—between the distributions. Note that we use quotation marks because, formally, the KL divergence is not a distance metric since it is not symmetric. Nevertheless, the variational objective is set to find a distribution $q^*_\phi$ such that

$$q^*_\phi(\underline{\theta}) = \min_\phi D_{KL}\left(q_\phi(\underline{\theta}) || \pi(\underline{\theta} \mid \underline{x})\right), \tag{S4}$$

where $D_{KL}$ is the KL divergence. Furthermore, we highlight that the KL divergence is a strictly positive number, i.e.,

$$D_{KL}\left(q_\phi(\underline{\theta}) || \pi(\underline{\theta} \mid \underline{x})\right) \geq 0, \tag{S5}$$

as this property will become important later on.

At first sight, Equation S4 does not improve the situation but only introduces further technical complications. After all, the definition of the KL divergence

$$D_{KL}\left(q_\phi(\underline{\theta}) || \pi(\underline{\theta} \mid \underline{x})\right) \equiv \int \cdots \int d^K\underline{\theta}\; q_\phi(\underline{\theta}) \ln \frac{q_\phi(\underline{\theta})}{\pi(\underline{\theta} \mid \underline{x})}, \tag{S6}$$

includes the posterior distribution $\pi(\underline{\theta} \mid \underline{x})$ we are trying to get around. However, let us manipulate Equation S6 to beat it to a more reasonable form. First, we can use the properties of the logarithms to write

$$D_{KL}\left(q_\phi(\underline{\theta}) || \pi(\underline{\theta} \mid \underline{x})\right) = \int d^K\underline{\theta}\; q_\phi(\underline{\theta}) \ln q_\phi(\underline{\theta}) - \int d^K\underline{\theta}\; q_\phi(\underline{\theta}) \ln \pi(\underline{\theta} \mid \underline{x}), \tag{S7}$$

where, for convenience, we write a single integration sign ($d^K\underline{\theta}$ still represents a multi-dimensional differential). For the second term in Equation S7, we can substitute the term inside the logarithm using Equation S2. This results in

$$\begin{aligned} D_{KL}\left(q_\phi(\underline{\theta}) || \pi(\underline{\theta} \mid \underline{x})\right) = &\int d^K\underline{\theta}\; q_\phi(\underline{\theta}) \ln q_\phi(\underline{\theta}) \\ &- \int d^K\underline{\theta}\; q_\phi(\underline{\theta}) \ln \left(\frac{\pi(\underline{x} \mid \underline{\theta})\pi(\underline{\theta})}{\pi(\underline{x})}\right). \end{aligned} \tag{S8}$$

Again, using the properties of logarithms, we can split Equation S8, obtaining

$$\begin{aligned} D_{KL}\left(q_\phi(\underline{\theta}) || \pi(\underline{\theta} \mid \underline{x})\right) = &\int d^K\underline{\theta}\; q_\phi(\underline{\theta}) \ln q_\phi(\underline{\theta}) \\ &- \int d^K\underline{\theta}\; q_\phi(\underline{\theta}) \ln \pi(\underline{x} \mid \underline{\theta}) \\ &- \int d^K\underline{\theta}\; q_\phi(\underline{\theta}) \ln \pi(\underline{\theta}) \\ &+ \int d^K\underline{\theta}\; q_\phi(\underline{\theta}) \ln \pi(\underline{x}). \end{aligned} \tag{S9}$$

It is convenient to write Equation S9 as

$$\begin{aligned} D_{KL}\left(q_\phi(\underline{\theta}) || \pi(\underline{\theta} \mid \underline{x})\right) = & \int d^K\underline{\theta}\; q_\phi(\underline{\theta}) \ln \frac{q_\phi(\underline{\theta})}{\pi(\underline{\theta})} \\ & - \int d^K\underline{\theta}\; q_\phi(\underline{\theta}) \ln \pi(\underline{x} \mid \underline{\theta}) \\ & + \ln \pi(\underline{x}) \int d^K\underline{\theta}\; q_\phi(\underline{\theta}), \end{aligned} \tag{S10}$$

where for the last term, we can take $\ln \pi(\underline{x})$ out of the integral since it does not depend on $\underline{\theta}$. Lastly, we utilize two properties:

1. The proposed approximate distribution must be normalized, i.e.,

$$\int d^K\underline{\theta}\; q_\phi(\underline{\theta}) = 1. \tag{S11}$$

2. The law of the unconscious statistician (LOTUS) establishes that for any probability density function, it must be true that

$$\int d^K\underline{\theta}\; q_\phi(\underline{\theta}) f(\underline{\theta}) = \left\langle f(\underline{\theta}) \right\rangle_{q_\phi}, \tag{S12}$$

   where $\langle \cdot \rangle_{q_\phi}$ is the expected value over the $q_\phi$ distribution.

Using these two properties, the positivity constraint on the KL divergence in Equation S5, and the definition of the KL divergence in Equation S6 we can rewrite Equation S10 as

$$D_{KL}\left(q_\phi(\underline{\theta}) || \pi(\underline{\theta})\right) - \left\langle \ln \pi(\underline{x} \mid \underline{\theta}) \right\rangle_{q_\phi} \geq - \ln \pi(\underline{x}). \tag{S13}$$

Multiplying by a minus one, we have the functional form of the so-called evidence lower bound (ELBO) Kingma and Welling [9],

$$\underbrace{\ln \pi(\underline{x})}_{\text{log evidence}} \geq \underbrace{\left\langle \ln \pi(\underline{x} \mid \underline{\theta}) \right\rangle_{q_\phi} - D_{KL}\left(q_\phi(\underline{\theta}) || \pi(\underline{\theta})\right)}_{\text{ELBO}}. \tag{S14}$$

Let us recapitulate where we are. We started by presenting the challenge of working with Bayes' theorem, as it requires a high-dimensional integral of the form in Equation S3. As an alternative, variational inference posits to approximate the posterior distribution $\pi(\underline{\theta} \mid \underline{x})$ with a parametric distribution $q_\phi(\underline{\theta})$. By minimizing the KL divergence between these distributions, we arrive at the result in Equation S14, where the left-hand side—the log marginalized likelihood or log evidence—we cannot compute for technical/computational reasons. However, the right-hand side is composed of things we can easily evaluate. We can easily evaluate the log-likelihood $\ln \pi(\underline{x} \mid \underline{\theta})$ and the KL divergence between our proposed approximate distribution $q_\phi(\underline{\theta})$ and the prior distribution $\pi(\underline{\theta})$. Moreover, we can compute the gradients of these functions with respect to the parameters of our proposed distribution. This last point implies that we can change the parameters of the proposed distribution to maximize the

ELBO. And, although we cannot compute the left-hand side of Equation S14, we know that however large we make the ELBO, it will always be smaller than (or equal) the log-marginal likelihood. Therefore, the larger we can make the ELBO by modifying the parameters $\phi$, the closer it gets to the log-marginal likelihood, and, as a consequence, the better our proposed distribution $q_\phi(\underline{\theta})$ gets to the true posterior distribution $\pi(\underline{\theta} \mid \underline{x})$.

In this sense, variational inference turns the intractable numerical integration problem to an optimization routine, for which there are several algorithms available.

#### ADVI algorithm

To maximize the right-hand side of Equation S14, the Automatic Differentiation Variational Inference (ADVI) algorithm developed in[7] takes advantage of advances in probabilistic programming languages to generate a robust method to perform this optimization. Without going into the details of the algorithm implementation, for our purposes, it suffices to say that we define our joint distribution $\pi(\underline{\theta}, \underline{x})$ as the product defined in Equation S1. ADVI then proposes an approximate variational distribution $q_\phi$ that can either be a multivariate Normal distribution with a diagonal covariance matrix, i.e.,

$$\phi = (\underline{\mu}, \underline{\underline{D}}), \tag{S15}$$

where $\underline{\underline{D}}$ is the identity matrix, with the diagonal elements given by the vector of variances $\underline{\sigma}^2$ for each variable or a full-rank multivariate Normal distribution

$$\phi = (\underline{\mu}, \underline{\underline{\Sigma}}). \tag{S16}$$

Then, the parameters are initialized in some value $\phi_o$. These parameters are iteratively updated by computing the gradient of the ELBO (right-hand side of Equation S14), hereafter defined as $\mathcal{L}$, with respect to the parameters,

$$\nabla_\phi \mathcal{L} = \nabla_{\underline{\mu}} \mathcal{L} + \nabla_{\underline{\sigma}} \mathcal{L}, \tag{S17}$$

and then computing

$$\phi_{t+1} = \phi_t + \eta \nabla_\phi \mathcal{L},$$

where $\eta$ defines the step size.

This short explanation behind the ADVI algorithm is intended only to gain intuition on how the optimal variational distribution $q_\phi$ be computed. There are many nuances in the implementation of the ADVI algorithm. We invite the reader to look at the original reference for further details.

#### Defining the Bayesian model

In the main text, we specify the inference problem we must solve as being of the form

$$\pi(\underline{s}^M, \underline{\bar{s}}_T, \underline{\underline{F}} \mid \underline{\underline{R}}) \propto \pi(\underline{\underline{R}} \mid \underline{s}^M, \underline{\bar{s}}_T, \underline{\underline{F}})\pi(\underline{s}^M, \underline{\bar{s}}_T, \underline{\underline{F}}). \tag{S18}$$

Here, we briefly define the missing nuisance parameters. Let

$$\underline{\bar{s}}_T = (\bar{s}_1, \bar{s}_2, \ldots, \bar{s}_{T-1})^\dagger, \tag{S19}$$

be the vector containing the $T-1$ population mean fitness we compute from the $T$ time points where measurements were taken. We have $T-1$ since the value of any $\bar{s}_t$ requires cycle numbers $t$ and $t+1$. Furthermore, let the matrix $\underline{\underline{F}}$ be a $T \times B$ matrix containing all frequency values. As with Equation 12 in the main text, we can split $\underline{\underline{F}}$ into two matrices of the form

$$\underline{\underline{F}} = \left[\underline{\underline{F}}^N \; \underline{\underline{F}}^M\right], \tag{S20}$$

to separate the corresponding neutral and non-neutral barcode frequencies.

Let us now define each of the terms in Equation 18 described in Section  of the main text. The following sections will specify the functional form each of these terms takes.

##### Frequency uncertainty $\pi(\underline{\underline{F}} \mid \underline{\underline{R}})$

We begin with the probability of the frequency values given the raw barcode reads. The first assumption is that the inference of the frequency values for time $t$ is independent of any other time. Therefore, we can write the joint probability distribution as a product of independent distributions of the form

$$\pi(\underline{\underline{F}} \mid \underline{\underline{R}}) = \prod_{t=1}^{T} \pi(\underline{f}_t \mid \underline{r}_t), \tag{S21}$$

where $\underline{f}_t$ and $\underline{r}_t$ are the $t$-th row of the matrix containing all of the measurements for time $t$. We imagine that when the barcode reads are obtained via sequencing, the quantified number of reads is a Poisson sample from the "true" underlying number of barcodes within the pool. This translates to assuming that the number of reads for each barcode at any time point $r_t^{(b)}$ is an independent Poisson random variable, i.e.,

$$r_t^{(b)} \sim \text{Poiss}(\lambda_t^{(b)}), \tag{S22}$$

where the symbol "$\sim$" is read "distributed as." Furthermore, for a Poisson distribution, we have that

$$\lambda_t^{(b)} = \left\langle r_t^{(b)} \right\rangle = \left\langle \left( r_t^{(b)} - \left\langle r_t^{(b)} \right\rangle \right)^2 \right\rangle, \tag{S23}$$

where $\langle \cdot \rangle$ is the expected value. In other words the Poisson parameter is equal to the mean and variance of the distribution. The Poisson distribution has the convenient property that for two Poisson distributed random variables $X \sim \text{Poiss}(\lambda_x)$ and $Y \sim \text{Poiss}(\lambda_y)$, we have that

$$Z \equiv X + Y \sim \text{Poiss}(\lambda_x + \lambda_y). \tag{S24}$$

This additivity allows us to write the total number of reads at time $t$ $n_t$ also as a Poisson-distributed random variable of the form

$$n_t \sim \text{Poiss}\left(\sum_{b=1}^{B} \lambda_t^{(b)}\right), \tag{S25}$$

where the sum is taken over all $B$ barcodes.

If the total number of reads is given by Equation S25, the array with the number of reads for each barcode at time $t$, $\underline{r}_t$ is then distributed as

$$\underline{r}_t \sim \text{Multinomial}(n_t, \underline{f}_t), \tag{S26}$$

where each of the $B$ entries of the frequency vector $\underline{f}_t$ is a function of the $\underline{\lambda}_t$ vector, given by

$$f_t^{(b)} \equiv f_t^{(b)}(\underline{\lambda}_t) = \frac{\lambda_t^{(b)}}{\sum_{b'=1}^{B} \lambda_t^{(b')}}. \tag{S27}$$

In other words, we can think of the $B$ barcode counts as independent Poisson samples or as a single multinomial draw with a random number of total draws, $n_t$, and the frequency vector $\underline{f}_t$ we are interested in. Notice that Equation S27 is a deterministic function that connects the Poisson parameters to the frequencies. Therefore, we have the equivalence that

$$\pi(\underline{f}_t \mid \underline{r}_t) = \pi(\underline{\lambda}_t \mid \underline{r}_t), \tag{S28}$$

meaning that the uncertainty comes from the $\underline{\lambda}_t$ vector. By Bayes theorem, we therefore write

$$\pi(\underline{\lambda}_t \mid n_t, \underline{r}_t) \propto \pi(n_t, \underline{r}_t \mid \underline{\lambda}_t)\pi(\underline{\lambda}_t), \tag{S29}$$

where we explicitly include the dependence on $n_t$. This does not affect the distribution or brings more uncertainty because $\underline{r}_t$ already contains all the information to compute $n_t$ since

$$n_t = \sum_{b=1}^{B} r_t^{(b)}. \tag{S30}$$

But adding the variable allows us to factorize Equation S29 as

$$\pi(\underline{\lambda}_t \mid n_t, \underline{r}_t) \propto \pi(\underline{r}_t \mid n_t, \underline{\lambda}_t)\pi(n_t \mid \underline{\lambda}_t)\pi(\underline{\lambda}_t) \tag{S31}$$

We then have

$$\underline{r}_t \mid n_t, \underline{\lambda}_t \sim \text{Multinomial}(n_t, \underline{f}_t(\underline{\lambda}_t)). \tag{S32}$$

Furthermore, we have

$$n_t \mid \underline{\lambda}_t \sim \text{Poiss}\left(\sum_{b=1}^{B} \lambda_t^{(b)}\right).$$

{#eq=freq_n_bayes} Finally, for our prior $\pi(\underline{\lambda}_t)$, we first assume each parameter is independent, i.e.,

$$\pi(\underline{\lambda}_t) = \prod_{b=1}^{B} \pi(\lambda_t^{(b)}).$$

A reasonable prior for each $\lambda_t^{(b)}$ representing the expected number of reads for barcode $b$ should span several orders of magnitude. Furthermore, we assume that no barcode in the dataset ever goes extinct. Thus, no frequency can equal zero, facilitating the computation of the log frequency ratios needed to infer the relative fitness. The log-normal distribution satisfies these constraints; therefore, for the prior, we assume

$$\lambda_t^{(b)} \sim \log\mathcal{N}(\mu_{\lambda_t^{(b)}}, \sigma_{\lambda_t^{(b)}}), \tag{S33}$$

with $\mu_{\lambda_t^{(b)}}, \sigma_{\lambda_t^{(b)}}$ as the user-defined parameters that characterize the prior distribution.

#### Summary

Putting all the pieces developed in this section together gives a term for our inference of the form

$$\pi(\underline{\underline{F}} \mid \underline{\underline{R}}) \propto \prod_{t=1}^{T} \left\{ \pi(\underline{r}_t \mid n_t, \underline{\lambda}_t) \pi(n_t \mid \underline{\lambda}_t) \left[ \prod_{b=1}^{B} \pi(\lambda_t^{(b)}) \right] \right\} \tag{S34}$$

where

$$\underline{r}_t \mid n_t, \underline{\lambda}_t \sim \text{Multinomial}(n_t, \underline{f}_t(\underline{\lambda}_t)), \tag{S35}$$

$$n_t \mid \underline{\lambda}_t \sim \text{Poiss}\left( \sum_{b=1}^{B} \lambda_t^{(b)} \right). \tag{S36}$$

and

$$\lambda_t^{(b)} \sim \log\mathcal{N}(\mu_{\lambda_t^{(b)}}, \sigma_{\lambda_t^{(b)}}), \tag{S37}$$

#### Population mean fitness uncertainty $\pi(\underline{\bar{s}}_T \mid \underline{\underline{F}}, \underline{\underline{R}})$

Next, we turn our attention to the problem of determining the population mean fitnesses $\underline{\bar{s}}_T$. First, we notice that our fitness model in Equation 3 does not include the value of the raw reads. They enter the calculation indirectly through the inference of the frequency values we developed in Section . This means that we can remove the conditioning of the value of $\underline{\bar{s}}_T$ on the number of reads, obtaining a simpler probability function

$$\pi(\underline{\bar{s}}_T \mid \underline{\underline{F}}, \underline{\underline{R}}) = \pi(\underline{\bar{s}}_T \mid \underline{\underline{F}}). \tag{S38}$$

Moreover, our fitness model does not directly explain how the population mean fitness evolves over time. In other words, our model cannot explicitly compute the population mean fitness at time $t+1$ from the information we have about time $t$. Given this model limitation, we are led to assume that we must infer each $\bar{s}_t$ independently. Expressing this for our inference results in

$$\pi(\underline{\bar{s}}_T \mid \underline{\underline{F}}) = \prod_{t=1}^{T-1} \pi(\bar{s}_t \mid \underline{f}_t, \underline{f}_{t+1}), \tag{S39}$$

where we split our matrix $\underline{\underline{F}}$ for each time point and only kept the conditioning on the relevant frequencies needed to compute the mean fitness at time $t$.

Although our fitness model in Equation 3 also includes the relative fitness $s^{(m)}$, to infer the population mean fitness we only utilize data from the neutral lineages that, by definition, have a relative fitness $s^{(n)} = 0$. Therefore, the conditioning on Equation S39 can be further simplified by only keeping the frequencies of the neutral lineages, i.e.,

$$\pi(\bar{s}_t \mid \underline{f}_t, \underline{f}_{t+1}) = \pi(\bar{s}_t \mid \underline{f}_t^N, \underline{f}_{t+1}^N). \tag{S40}$$

Recall that in Section  we emphasized that the frequencies $f_t^{(n)}$ do not represent the true frequency of a particular lineage in the population but rather a "normalized number of cells." Therefore, it is safe to assume each of the $N$ neutral lineages' frequencies is changing independently. The correlation of how increasing the frequency of one lineage will decrease the frequency of others is already captured in the model presented in Section . Thus, we write

$$\pi(\bar{s}_t \mid \underline{f}_t^N, \underline{f}_{t+1}^N) = \prod_{n=1}^{N} \pi(\bar{s}_t \mid f_t^{(n)}, f_{t+1}^{(n)}). \tag{S41}$$

Now, we can focus on one of the terms on the right-hand side of Equation S41. Writing Bayes theorem results in

$$\pi(\bar{s}_t \mid f_t^{(n)}, f_{t+1}^{(n)}) \propto \pi(f_t^{(n)}, f_{t+1}^{(n)} \mid \bar{s}_t)\pi(\bar{s}_t). \tag{S42}$$

Notice the likelihood defines the joint distribution of neutral barcode frequencies conditioned on the population mean fitness. However, rewriting our fitness model in Equation 3 for a neutral lineage to leave frequencies on one side and fitness on the other results in

$$\frac{f_{t+1}^{(n)}}{f_t^{(n)}} = \mathrm{e}^{-\bar{s}_t \tau}. \tag{S43}$$

Equation S43 implies that our fitness model only relates **the ratio** of frequencies and not the individual values. To get around this complication, we define

$$\gamma_t^{(b)} \equiv \frac{f_{t+1}^{(b)}}{f_t^{(b)}}, \tag{S44}$$

as the ratio of frequencies between two adjacent time points for any barcode $b$. This allows us to rewrite the joint distribution $\pi(f_t^{(n)}, f_{t+1}^{(n)} \mid \bar{s}_t)$ as

$$\pi(f_t^{(n)}, f_{t+1}^{(n)} \mid \bar{s}_t) = \pi(f_t^{(n)}, \gamma_t^{(n)} \mid \bar{s}_t). \tag{S45}$$

Let us rephrase this subtle but necessary change of variables since it is a key part of the inference problem: our series of independence assumptions lead us to Equation S42 that relates the value of the population mean fitness $\bar{s}_t$ to the frequency of a neutral barcode at times $t$ and $t+1$. However, as shown in Equation S43, our model functionally relates the ratio of frequencies—that we defined as $\gamma_t^{(n)}$—and not the independent frequencies to the mean fitness. Therefore, instead of writing for the likelihood the joint distribution of the frequency values at times $t$ and $t+1$ conditioned on the mean fitness, we write the joint distribution of the barcode frequency at time $t$ and the ratio of the frequencies. These **must be** equivalent joint distributions since there is a one-to-one mapping between $\gamma_t^{(n)}$ and $f_{t+1}^{(n)}$ for a given value of $f_t^{(n)}$. Another way to phrase this is to say that knowing the frequency at time $t$ and at time $t+1$ provides the same amount of information as knowing the frequency at time $t$ and the ratio of the frequencies. This is because if we want to obtain $f_{t+1}^{(n)}$ given this information, we simply compute

$$f_{t+1}^{(n)} = \gamma_t^{(n)} f_t^{(n)}. \tag{S46}$$

The real advantage of rewriting the joint distribution as in Equation S45 comes from splitting this joint distribution as a product of conditional distributions of the form

$$\pi(f_t^{(n)}, \gamma_t^{(n)} \mid \bar{s}_t) = \pi(f_t^{(n)} \mid \gamma_t^{(n)}, \bar{s}_t)\pi(\gamma_t^{(n)} \mid \bar{s}_t). \tag{S47}$$

Written in this form, we can finally propose a probabilistic model for how the mean fitness relates to the frequency ratios we determine in our experiments. The second term on the right-hand side of Equation S47 relates how the determined frequency ratio $\gamma_t^{(b)}$ relates to the mean fitness $\bar{s}_t$. From Equation S43 and Equation S44, we can write

$$\ln \gamma_t^{(n)} = -\bar{s}_t + \varepsilon_t^{(n)}, \tag{S48}$$

where, for simplicity, we set $\tau = 1$. Note that we added an extra term, $\varepsilon_t^{(n)}$, characterizing the deviations of the measurements from the theoretical model. We assume these errors are normally distributed with mean zero and some standard deviation $\sigma_t$, implying that

$$\ln \gamma_t^{(n)} \mid \bar{s}_t, \sigma_t \sim \mathcal{N}\left(-\bar{s}_t, \sigma_t\right), \tag{S49}$$

where we include the nuisance parameter $\sigma_t$ to be determined. If we assume the log frequency ratio is normally distributed, this implies the frequency ratio itself is distributed log-normal. This means that

$$\gamma_t^{(n)} \mid \bar{s}_t, \sigma_t \sim \log\mathcal{N}\left(-\bar{s}_t, \sigma_t\right). \tag{S50}$$

Having added the nuisance parameter $\sigma_t$ implies that we must update Equation S42 to

$$\pi(\bar{s}_t, \sigma_t \mid f_t^{(n)}, f_{t+1}^{(n)}) \propto \pi(f_t^{(n)}, \gamma_t^{(n)} \mid \bar{s}_t, \sigma_t)\pi(\bar{s}_t)\pi(\sigma_t), \tag{S51}$$

where we assume the prior for each parameter is independent, i.e.,

$$\pi(\bar{s}_t, \sigma_t) = \pi(\bar{s}_t)\pi(\sigma_t). \tag{S52}$$

For numerical stability, we will select weakly-informative priors for both of these parameters. In the case of the nuisance parameter $\sigma_t$, the prior must be restricted to positive values only, since standard deviations cannot be negative.

For the first term on the right-hand side of Equation S47, $\pi(f_t^{(n)} \mid \gamma_t^{(n)}, \bar{s}_t)$, we remove the conditioning on the population mean fitness since it does not add any information on top of what the frequency ratio $\gamma_t^{(n)}$ already gives. Therefore, we have

$$\pi(f_t^{(n)} \mid \gamma_t^{(n)}, \bar{s}_t) = \pi(f_t^{(n)} \mid \gamma_t^{(n)}). \tag{S53}$$

The right-hand side of Equation S53 asks us to compute the probability of observing a frequency value $f_t^{(n)}$ given that we get to observe the ratio $\gamma_t^{(n)}$. If the ratio happened to be $\gamma_t^{(n)} = 2$, we could have $f_{t+1}^{(n)} = 1$ and $f_{t+1}^{(n)} = 0.5$, for example. Although, it would be equally likely that $f_{t+1}^{(n)} = 0.6$ and $f_{t+1}^{(n)} = 0.3$ or $f_{t+1}^{(n)} = 0.1$ and $f_{t+1}^{(n)} = 0.05$ for that matter. If we only get to observe the frequency ratio $\gamma_t^{(n)}$, we know that the numerator $f_{t+1}^{(n)}$ can only take values between zero and one, all of them being equally likely given only the information on the ratio. As a consequence, the value of the frequency in the denominator $f_t^{(n)}$ is restricted to fall in the range

$$f_t^{(n)} \in \left(0, \frac{1}{\gamma_t^{(n)}}\right]. \tag{S54}$$

A priori, we do not have any reason to favor any value over any other, therefore it is natural to write

$$f_t^{(n)} \mid \gamma_t^{(n)} \sim \text{Uniform}\left(0, \frac{1}{\gamma_t^{(n)}}\right). \tag{S55}$$

#### Summary

Putting all the pieces we have developed in this section together results in an inference for the population mean fitness values of the form

$$\pi(\underline{\bar{s}}_T, \underline{\sigma}_T \mid \underline{\underline{F}}) \propto \prod_{t=1}^{T-1} \left\{ \prod_{n=1}^{N} \left[ \pi(f_t^{(n)} \mid \gamma_t^{(n)}) \pi(\gamma_t^{(n)} \mid \bar{s}_t, \sigma_t) \right] \pi(\bar{s}_t) \pi(\sigma_t) \right\}, \tag{S56}$$

where we have

$$f_t^{(n)} \mid \gamma_t^{(n)} \sim \text{Uniform}\left(0, \frac{1}{\gamma_t^{(n)}}\right), \tag{S57}$$

$$\gamma_t^{(n)} \mid \bar{s}_t, \sigma_t \sim \log\mathcal{N}(\bar{s}_t, \sigma_t), \tag{S58}$$

$$\bar{s}_t \sim \mathcal{N}(0, \sigma_{\bar{s}_t}), \tag{S59}$$

and

$$\sigma_t \sim \log\mathcal{N}(\mu_{\sigma_t}, \sigma_{\sigma_t}), \tag{S60}$$

where $\sigma_{\bar{s}_t}$, $\mu_{\sigma_t}$, and $\sigma_{\sigma_t}$ are user-defined parameters.

##### Mutant relative fitness uncertainty $\pi(\underline{s}^M \mid \underline{\bar{s}}_T, \underline{\underline{F}}, \underline{\underline{R}})$

The last piece of our inference is the piece that we care about the most: the probability distribution of all the mutants' relative fitness, given the inferred population mean fitness and the frequencies. First, we assume that all fitness values are independent of each other. This allows us to write

$$\pi(\underline{s}^M \mid \underline{\bar{s}}_T, \underline{\underline{F}}, \underline{\underline{R}}) = \prod_{m=1}^{M} \pi(s^{(m)} \mid \underline{\bar{s}}_T, \underline{\underline{F}}, \underline{\underline{R}}). \tag{S61}$$

Furthermore, as was the case with the population mean fitness, our fitness model relates frequencies, not raw reads. Moreover, the fitness value of mutant $m$ only depends on the frequencies of such mutant. Therefore, we can simplify the conditioning to

$$\pi(s^{(m)} \mid \underline{\bar{s}}_T, \underline{\underline{F}}, \underline{\underline{R}}) = \pi(s^{(m)} \mid \underline{\bar{s}}_T, \underline{f}^{(m)}), \tag{S62}$$

where

$$\underline{f}^{(m)} = (f_0^{(m)}, f_1^{(m)}, \ldots, f_T^{(m)})^\dagger, \tag{S63}$$

is the vector containing the frequency time series for mutant $m$. Writing Bayes' theorem for the right-hand side of Equation S62 results in

$$\pi(s^{(m)} \mid \underline{\bar{s}}_T, \underline{f}^{(m)}) \propto \pi(\underline{f}^{(m)} \mid \underline{\bar{s}}_T, s^{(m)}) \pi(s^{(m)} \mid \underline{\bar{s}}_T). \tag{S64}$$

Notice the conditioning on the mean fitness values $\underline{\bar{s}}_T$ is not inverted since we already inferred these values.

Following the logic used in Section , let us define

$$\underline{\gamma}^{(m)} = (\gamma_0^{(m)}, \gamma_1^{(m)}, \ldots, \gamma_{T-1}^{m})^\dagger, \tag{S65}$$

where each entry $\gamma_t^{(m)}$ is defined by Equation S44. In the same way we rewrote the joint distribution between two adjacent time point frequencies to the joint distribution between one of the frequencies and the ratio of both frequencies in Equation S45, we can rewrite the joint distribution of the frequency time series for mutant $m$ as

$$\pi(\underline{f}^{(m)} \mid \underline{\bar{s}}_T, s^{(m)}) = \pi(f_0^{(m)}, \underline{\gamma}^{(m)} \mid \underline{\bar{s}}_T, s^{(m)}). \tag{S66}$$

One can think about Equation S66 as saying that knowing the individual frequencies at each time point contain equivalent information as knowing the initial frequency and the subsequent ratios of frequencies. This is because if we want to know the value of $f_1^{(m)}$ given the ratios, we only need to compute

$$f_1^{(m)} = \gamma_0^{(m)} f_0^{(m)}. \tag{S67}$$

Moreover, if we want to know $f_2^{(m)}$, we have

$$f_2^{(m)} = \gamma_1^{(m)} f_1^{(m)} = \gamma_1^{(m)} \left(\gamma_0^{(m)} f_0^{(m)}\right), \tag{S68}$$

and so on. We can then write the joint distribution on the right-hand side of Equation S66 as a product of conditional distributions of the form

$$
\begin{aligned}
\pi(f_0^{(m)}, \underline{\gamma}^{(m)} \mid \underline{\bar{s}}_T, s^{(m)}) =& \pi(f_0^{(m)} \mid \gamma_0^{(m)}, \ldots, \gamma_{T-1}^{(m)}, \underline{\bar{s}}_T, s^{(m)}) \times \\
& \pi(\gamma_0^{(m)} \mid \gamma_1^{(m)}, \ldots, \gamma_{T-1}^{(m)}, \underline{\bar{s}}_T, s^{(m)}) \times \\
& \pi(\gamma_1^{(m)} \mid \gamma_2^{(m)}, \ldots, \gamma_{T-1}^{(m)}, \underline{\bar{s}}_T, s^{(m)}) \times \\
& \vdots \\
& \pi(\gamma_{T-2}^{(m)} \mid \gamma_{T-1}^{(m)}, \underline{\bar{s}}_T, s^{(m)}) \times \\
& \pi(\gamma_{T-1}^{(m)} \mid \underline{\bar{s}}_T, s^{(m)}).
\end{aligned}
\tag{S69}
$$

Writing the fitness model in Equation 3 as

$$
\gamma_t^{(m)} = \frac{f_{t+1}^{(m)}}{f_t^{(m)}} = \mathrm{e}^{(s^{(m)} - s_t)\tau},
$$

reveals that the value of each of the ratios $\gamma_t^{(m)}$ only depends on the corresponding fitness value $\bar{s}_t$ and the relative fitness $s^{(m)}$. Therefore, we can remove most of the conditioning on the right-hand side of Equation S69, resulting in a much simpler joint distribution of the form

$$
\begin{aligned}
\pi(f_0^{(m)}, \underline{\gamma}^{(m)} \mid \underline{\bar{s}}_T, s^{(m)}) =& \pi(f_0^{(m)} \mid \gamma_0^{(m)}) \times \\
& \pi(\gamma_0^{(m)} \mid \bar{s}_0, s^{(m)}) \times \\
& \pi(\gamma_1^{(m)} \mid \bar{s}_1, s^{(m)}) \times \\
& \vdots \\
& \pi(\gamma_{T-2}^{(m)} \mid \bar{s}_{T-2}, s^{(m)}) \times \\
& \pi(\gamma_{T-1}^{(m)} \mid \bar{s}_{T-1}, s^{(m)}),
\end{aligned}
\tag{S70}
$$

where for the first term on the right-hand side of Equation S70 we apply the same logic as in Equation S53 to remove all other dependencies. We emphasize that although Equation S70 looks like a series of independent inferences, the value of the relative fitness $s^{(m)}$ is shared among all of them. This means that the parameter is not inferred individually for each time point, resulting in different estimates of the parameter, but each time point contributes independently to the inference of a single estimate of $s^{(m)}$.

Using equivalent arguments to those in Section , we assume

$$
f_0^{(m)} \mid \gamma_0^{(m)} \sim \mathrm{Uniform}\left(0, \frac{1}{\gamma_0^{(m)}}\right),
$$

and

$$
\gamma_t^{(m)} \mid \bar{s}_t, s^{(m)}, \sigma^{(m)} \sim \log\mathcal{N}\left(s^{(m)} - \bar{s}_t, \sigma^{(m)}\right),
\tag{S71}
$$

where we add the nuisance parameter $\sigma^{(m)}$ to the inference. Notice that this parameter is not indexed by time. This means that we assume the deviations from the theoretical prediction do not depend on time, but only on the mutant. Adding the nuisance parameter demands us to update Equation S64 to

$$\pi(s^{(m)}, \sigma^{(m)} \mid \underline{\bar{s}}_T, \underline{f}^{(m)}) \propto \pi(\underline{f}^{(m)} \mid \underline{\bar{s}}_T, s^{(m)}, \sigma^{(m)})\pi(s^{(m)})\pi(\sigma^{(m)}), \tag{S72}$$

where we assume independent priors for both parameters. We also removed the conditioning on the values of the mean fitness as knowing such values does not change our prior information about the possible range of values that the parameters can take. As with the priors on Section , we will assign weakly-informative priors to these parameters.

#### Summary

With all pieces in place, we write the full inference of the relative fitness values as

$$\pi(\underline{s}^M, \underline{\sigma}^M \mid \underline{\bar{s}}_T, \underline{\underline{F}}) \propto \prod_{m=1}^{M} \left\{ \pi(f_0^{(m)} \mid \gamma_0^{(m)}) \prod_{t=0}^{T-1} \left[ \pi(\gamma_t^{(m)} \mid \bar{s}_t, s^{(m)}, \sigma^{(m)}) \right] \pi(s^{(m)})\pi(\sigma^{(m)}) \right\}, \tag{S73}$$

where

$$f_0^{(m)} \mid \gamma_0^{(m)} \sim \text{Uniform}\left(0, \frac{1}{\gamma_0^{(m)}}\right), \tag{S74}$$

$$\gamma_t^{(m)} \mid \bar{s}_t, s^{(m)}, \sigma^{(m)} \sim \log\mathcal{N}\left(s^{(m)} - \bar{s}_t, \sigma^{(m)}\right), \tag{S75}$$

$$s^{(m)} \sim \mathcal{N}(0, \sigma_{s^{(m)}}), \tag{S76}$$

and

$$\sigma^{(m)} \sim \log\mathcal{N}(\mu_{\sigma^{(m)}}, \sigma_{\sigma^{(m)}}), \tag{S77}$$

where $\sigma_{s^{(m)}}$, $\mu_{\sigma^{(m)}}$, and $\sigma_{\sigma^{(m)}}$ are user-defined parameters.

#### Hierarchical models for multiple experimental replicates

As detailed in Section  of the main text, we define a Bayesian hierarchical model to analyze data from multiple experimental replicates. The implementation requires only slightly modifying the base model detailed in the previous sections. The hierarchical model defines a hyper-fitness parameter $\theta^{(m)}$ for every non-neutral barcode. We can thus collect all of the $M$ hyperparameters in an array of the form

$$\underline{\theta}^M = (\theta^{(1)}, \ldots, \theta^{(M)})^\dagger. \tag{S78}$$

Our data now consists of a series of matrices $\underline{\underline{R}}_{[j]}$, where the subindex $[j]$ refers to the $j$-th experimental replicate. These matrices need not have the same number of rows, as the time points measured for each replicate can vary. The statistical model we must define is then of the form

$$\pi(\underline{\theta}^M, \{\underline{s}^M_{[j]}\}, \{\bar{s}_{T[j]}\}, \{\underline{\underline{F}}_{[j]}\} \mid \{\underline{\underline{R}}_{[j]}\}) \propto \pi(\{\underline{\underline{R}}_{[j]}\} \mid \underline{\theta}^M, \{\underline{s}^M_{[j]}\}, \{\bar{s}_{T[j]}\}, \{\underline{\underline{F}}_{[j]}\}) \times \pi(\underline{\theta}^M, \{\underline{s}^M_{[j]}\}, \{\bar{s}_{T[j]}\}, \{\underline{\underline{F}}_{[j]}\}) \tag{S79}$$

where the parameters within curly braces with subindex $[j]$ indicate one set of parameters per experimental replicate. For example,

$$\{\underline{s}^M_{[j]}\} = \{\underline{s}^M_{[1]}, \underline{s}^M_{[2]}, \ldots \underline{s}^M_{[E]}\}, \tag{S80}$$

where $E$ is the number of experimental replicates.

Given the dependencies between the variables, we can factorize Equation S79 to be of the form

$$\pi(\underline{\theta}^M, \{\underline{s}^M_{[j]}\}, \{\bar{s}_{T[j]}\}, \{\underline{\underline{F}}_{[j]}\} \mid \{\underline{\underline{R}}_{[j]}\}) = \pi(\underline{\theta}^M, \{\underline{s}^M_{[j]}\} \mid \{\bar{s}_{T[j]}\}, \{\underline{\underline{F}}_{[j]}\}) \times \pi(\{\bar{s}_{T[j]}\} \mid \{\underline{\underline{F}}_{[j]}\}) \times \pi(\{\underline{\underline{F}}_{[j]}\} \mid \{\underline{\underline{R}}_{[j]}\}) \tag{S81}$$

Furthermore, the hierarchical structure only connects the replicates via the relative fitness parameters. This means that the population mean fitness values and the frequencies can be independently inferred for each dataset. This allows us to rewrite the right-hand side of Equation S81 as

$$\pi(\underline{\theta}^M, \{\underline{s}^M_{[j]}\}, \{\bar{s}_{T[j]}\}, \{\underline{\underline{F}}_{[j]}\} \mid \{\underline{\underline{R}}_{[j]}\}) = \pi(\underline{\theta}^M, \{\underline{s}^M_{[j]}\} \mid \{\bar{s}_{T[j]}\}, \{\underline{\underline{F}}_{[j]}\}) \times \prod_{j=1}^{E} \left[\pi(\bar{s}_{T[j]} \mid \underline{\underline{F}}_{[j]}) \pi(\underline{\underline{F}}_{[j]} \mid \underline{\underline{R}}_{[j]})\right]. \tag{S82}$$

The terms inside the square brackets in Equation S82 take the same functional form as those derived in Section  and Section . Therefore, to implement the desired hierarchical model, we only need to focus on the first term on the right-hand side of Equation S82. A way to think about the structure of the hierarchical model is as follows: imagine each genotype as a "*true*" relative fitness value. However, every time we perform an experiment, small variations in the biotic and abiotic conditions—also known as batch effects—might result in small deviations from this value. We model this by defining a distribution for the hyper-fitness parameter—the ground truth we are interested in—and having each experimental replicate sample from this hyper-parameter distribution to determine the "*local*" fitness value. The wider the hyper-parameter distribution is the more variability between experimental replicates.

Writing Bayes' theorem for the first term in Equation S82 results in

$$\pi(\underline{\theta}^M, \{\underline{s}^M_{[j]}\} \mid \{\underline{\underline{F}}_{[j]}\}, \{\bar{s}_{T[j]}\}) \propto \pi(\{\underline{\underline{F}}_{[j]}\} \mid \underline{\theta}^M, \{\underline{s}^M_{[j]}\}, \{\bar{s}_{T[j]}\}) \pi(\underline{\theta}^M, \{\underline{s}^M_{[j]}\} \mid \{\bar{s}_{T[j]}\}), \tag{S83}$$

where we leave the conditioning on the population mean fitness as we did in Section . This expression can be simplified in two ways. First, the frequency values for each experimental replicate depend directly on the local fitness values and the corresponding population mean fitness, as the relationship between experimental replicates only occurs through the relative fitness values. Therefore, we can write

$$\pi(\underline{\theta}^M, \{\underline{s}^M_{[j]}\} \mid \{\underline{\underline{F}}_{[j]}\}, \{\bar{s}_{T[j]}\}) \propto \prod_{j=1}^{E} \left[\pi(\underline{\underline{F}}_{[j]} \mid \underline{s}^M_{[j]}, \bar{s}_{T[j]})\right] \pi(\underline{\theta}^M, \{\underline{s}^M_{[j]}\} \mid \{\bar{s}_{T[j]}\}). \tag{S84}$$

Second, the relationship between the hyper-fitness and the local fitness values allows us to write their joint distribution as a conditional distribution where local fitness values depend on the global hyper-fitness value, obtaining

$$\pi(\underline{\theta}^M, \{\underline{s}^M_{[j]}\} \mid \{\underline{\underline{F}}_{[j]}\}, \{\bar{s}_{T[j]}\}) \propto \prod_{j=1}^{E} \left[\pi(\underline{\underline{F}}_{[j]} \mid \underline{s}^M_{[j]}, \bar{s}_{T[j]}) \pi(\underline{s}^M_{[j]} \mid \underline{\theta}^M)\right] \pi(\underline{\theta}^M). \tag{S85}$$

Notice we removed the conditioning on the population mean fitness as our prior expectations of what the global hyper-fitness or local fitness value might be do not depend on these nuisance parameters.

The first term on the right-hand side of Equation S85 takes the same functional form as the one derived in Section . Therefore, all we are left with is to determine the functional forms for the hyper–prior $\pi(\underline{\theta}^M)$, and the conditional probability $\pi(\underline{s}^M_{[j]} \mid \underline{\theta}^M)$. In analogy to the assumptions used for the fitness values in Section , we define the value of each hyper-fitness as independent. This means that we have

$$\pi(\underline{\theta}^M) = \prod_{m=1}^{M} \pi(\theta^{(m)}). \tag{S86}$$

Furthermore, we assume this prior is of the form

$$\theta^{(m)} \sim \mathcal{N}(\mu_{\theta^{(m)}}, \sigma_{\theta^{(m)}}), \tag{S87}$$

where $\mu_{\theta^{(m)}}$ and $\sigma_{\theta^{(m)}}$ are user-defined parameters encoding the prior expectations on the fitness values.

For the conditional distribution $\pi(\underline{s}^M_{[j]} \mid \underline{\theta}^M)$, we use the so-called non-centered parametrization that avoids some of the intrinsic degeneracies associated with hierarchical models[13]. We invite the reader to check this excellent blog explaining the difficulties of working with hierarchical models. This non-centered parameterization implies that we introduce two nuisance parameters such that the local fitness $s^{(m)}_{[j]}$ is computed as

$$s^{(m)}_{[j]} = \theta^{(m)} + (\tau^{(m)}_{[j]} \times \xi^{(m)}_{[j]}), \tag{S88}$$

where $\theta^{(m)}$ is the corresponding genotype hyper-fitness value, $\xi^{(m)}_{[j]}$ is a standard normal random variable, i.e.,

$$\xi^{(m)}_{[j]} \sim \mathcal{N}(0, 1), \tag{S89}$$

that allows deviations from the hyper-fitness value to be either positive or negative, and $\tau_{[j]}^{(m)}$ is a strictly positive random variable that characterizes the deviation of the local fitness value from the global hyper-fitness. We assume

$$\tau_{[j]}^{(m)} \sim \log \mathcal{N}(\mu_{\tau_{[j]}^{(m)}}, \sigma_{\tau_{[j]}^{(m)}}) \tag{S90}$$

where $\mu_{\tau_{[j]}^{(m)}}$ and $\sigma_{\tau_{[j]}^{(m)}}$ are user-defined parameters capturing the expected magnitude of the batch effects.

#### Defining prior probabilities

One aspect commonly associated—in both positive and negative ways—to Bayesian analysis is the definition of prior probabilities. On the one hand, the naive textbook version of Bayesian analysis defines the prior as encoding the information we have about the inference in question before acquiring any data. This is the "ideal" use of priors that, whenever possible, should be implemented. On the other hand, for most practitioners of Bayesian statistics in the age of big data, the definition of prior becomes a tool to ensure the convergence of sampling algorithms such as MCMC[25]. However, for our particular problem, although we deal with large amounts of data (inferences can be made for $>$ 10K barcodes over multiple time points, resulting in $>$ 100K parameters), each barcode has very little data, as they are measured only once per time point over $<$ 10 growth-dilution cycles. Furthermore, it is incredibly challenging to understand the noise sources related to culturing conditions, DNA extraction, library preparation, etc., and encode them into reasonable prior distributions.

Empirically, our approach for this work defined the priors based solely on the neutral lineage data, as they represent the only repeated measurements of a single genotype in our experimental design. We acknowledge that defining the priors after observing the data might be considered an incoherent inference. However, as expressed by Gelman et al. [25]

> Incoherence is an unavoidable aspect of much real-world data analysis; and, indeed, one might argue that as scientists we learn the most from the anomalies and reassessments associated with episodes of incoherence.

With this in mind, we leave it to the reader to judge the selection of priors. Furthermore, the software package associated with this work, `BarBay.jl`, is written so that users can experiment with different prior selection criteria that fit their needs. We strongly advocate that statistics should not be done in a black-box fit-all tool mindset but rather as a formal way to encode the assumptions behind the analysis, subject to constructive criticism. With this philosophical baggage behind us, let us now focus on how the priors used for this work were selected.

#### Naive neutral lineage-based priors

For the base model presented in this work, the user-defined prior parameters include the following:

- Prior on population mean fitness (one per pair of adjacent time points)

$$\bar{s}_t \sim \mathcal{N}(\mu_{\bar{s}_t}, \sigma_{\bar{s}_t}). \tag{S91}$$

- Prior on standard deviation associated with neutral lineages likelihood function (one per pair of adjacent time points)

$$\sigma_t \sim \log\mathcal{N}(\mu_{\sigma_t}, \sigma_{\sigma_t}). \tag{S92}$$

- Prior on relative fitness (one per non-neutral barcode)

$$s^{(m)} \sim \mathcal{N}(\mu_{s^{(m)}}, \sigma_{s^{(m)}}). \tag{S93}$$

- Prior on standard deviation associated with non-neutral lineages likelihood function (one per non-neutral barcode)

$$\sigma^{(m)} \sim \log\mathcal{N}(\mu_{\sigma^{(m)}}, \sigma_{\sigma^{(m)}}) \tag{S94}$$

The `BarBay.jl` package includes a function `naive_prior` within the `stats` module. This function utilizes the data from the neutral lineages to determine some of the prior parameters to facilitate the inference algorithm's numerical convergence. In particular, it defines the population mean fitness parameter $\mu_{\bar{s}_t}$ as

$$\mu_{\bar{s}_t} = \frac{1}{N}\sum_{n=1}^{N} -\ln\left(\frac{r_{t+1}^{(n)}}{r_t^{(n)}}\right), \tag{S95}$$

where $N$ is the number of neutral lineages and $r_t^{(n)}$ is the number of neutral lineages. In other words, it defines the mean of the prior distribution as the mean of what one naively would compute from the neutral lineages, discarding cases where the ratio diverges because the denominator $r_t^{(n)} = 0$. For the variance parameter, we chose a value $\sigma_{\bar{s}_t} = 0.05$.

Furthermore, the `naive_prior` function defines the mean of the variance parameter as the standard deviation of the log frequency ratios for the neutral lineages, i.e.,

$$\mu_{\sigma_t} = \sqrt{\mathrm{Var}\left(\frac{r_{t+1}^{(n)}}{r_t^{(n)}}\right)}, \tag{S96}$$

where $\mathrm{Var}$ is the sample variance. This same value was utilized for the mean of the non-neutral barcode variance $\mu_{\sigma^{(m)}}$. While we assigne the corresponding variances to be $\sigma_{\sigma_t} = \sigma_{\sigma^{(m)}} = 1$.

#### Posterior predictive checks

Throughout the main text, we allude to the concept of posterior predictive checks as a formal way to assess the accuracy of our inference pipeline. Here, we explain the mechanics behind the computation of these credible regions, given the output of the inference.

Bayesian models encode what is known as a *generative model*. This statement means that in our definition of the likelihood function and the prior distribution, we, as modelers, propose a mathematical function that captures all relevant relationships between unobserved (latent) variables. Therefore, when these latent variables are input into the mathematical model, this function *generates* data that should be, in principle, indistinguishable from the real observations if the model is a good account of the underlying processes involved in the phenomena of interest. This generative model implies that once we run the inference process and update our posterior beliefs about the state of the latent variables, we can input back the inferred values to our model and generate synthetic data. Furthermore, we can repeat this process multiple times to compute the range where we expect to observe our data conditioned on the accuracy of the model.

For our specific scenario, recall that our objective is to infer the relative fitness of a non-neutral lineage $s^{(m)}$ along with nuisance parameters related to the population mean fitness at each point, $\underline{\bar{s}}_t$, and the barcode frequency time series $\underline{f}^{(m)}$. All these variables are related through our fitness model (see Section  in the main text)

$$f_{t+1}^{(m)} = f_t^{(m)} \mathrm{e}^{(s^{(m)} - s_t)\tau}. \tag{S97}$$

As we saw, it is convenient to rewrite Equation S97 as

$$\frac{1}{\tau} \ln \frac{f_{t+1}^{(m)}}{f_t^{(m)}} = (s^{(m)} - s_t). \tag{S98}$$

Written in this way, we separate the quantities we can compute from the experimental observations—the left-hand side of Equation S98 can be computed from the barcode reads—from the latent variables.

Although we perform the joint inference over all barcodes in the present work, let us focus on the inference task for a single barcode as if it were computed independently. For a non-neutral barcode, our task consists of computing the posterior probability

$$\pi(\theta \mid \underline{r}^{(m)}) = \pi(s^{(m)}, \sigma^{(m)}, \underline{s}_t, \underline{f}^{(m)} \mid \underline{r}^{(m)}), \tag{S99}$$

where $\theta$ represents all parameters to be inferred and $\underline{r}^{(m)}$ is the vector with the barcode raw counts time series. The list of parameters are

- $s^{(m)}$: The barcode's relative fitness.
- $\sigma^{(m)}$: A nuisance parameter used in the likelihood to generate the data. This captures the expected deviation from Equation S98

- $\underline{s}_t$: The vector with all population mean fitness for each pair of adjacent time points.
- $\underline{f}^{(m)}$: The vector with the barcode frequency time series.

Furthermore, let us define a naive estimate of the barcode frequency at time $t$ as

$$\hat{f}_t^{(m)} = \frac{r_t^{(m)}}{\sum_{b=1}^{B} r_t^{(b)}}. \tag{S100}$$

We can compute this quantity from the data by normalizing the raw barcode counts by the sum of all barcode counts. Furthermore, we can compute a naive estimate of the log frequency ratio from the raw barcode counts as

$$\ln \hat{\gamma}_t^{(m)} = \ln \frac{\hat{f}_{t+1}^{(m)}}{\hat{f}_t^{(m)}} \tag{S101}$$

In our generative model, we assumed

$$\ln \gamma_t^{(m)} \mid \theta \sim \mathcal{N}(s^{(m)} - s_t, \sigma^{(m)}). \tag{S102}$$

This implies that once we determine the posterior distribution of our parameters, we can generate synthetic values of $\ln \gamma_t^{(m)}$ that we can then compare with the values obtained from applying Equation S101 and Equation S101 to the raw data.

In practice, to compute the posterior predictive checks, we generate multiple samples from the posterior distribution $\pi(\theta \mid \underline{r}^{(m)})$

$$\underline{\theta} = (\theta_1, \theta_2, \ldots, \theta_N). \tag{S103}$$

With these samples in hand, the `BarBay.jl` package includes the function `logfreq_ratio_bc_ppc` for non-neutral barcodes that uses this set of posterior parameter samples to generate samples from the distribution defined in Equation S102. For a large-enough number of samples, we can then compute the desired percentiles—5, 68, and 95 percentiles in all figures in the main text—that are equivalent to the corresponding credible regions. In other words, the range of values of $\ln \gamma_t^{(m)}$ generated by this bootstrap process can be used to compute the region where we expect to find our raw estimates $\ln \hat{\gamma}_t^{(m)}$ with the desired probability. The package `BarBay.jl` includes an equivalent function, `logfreq_ratio_popmean_ppc`, for neutral lineages.

#### Logistic growth simulation

In this section, we explain the simulations used to assess the validity of our inference pipeline. Let us begin by assuming that, since the strains are grown for two full days in the experiment, having left behind the exponential phase for almost an entire day, a simple exponential growth of the form

$$\frac{dn_i}{dt} = \lambda_i n_i, \tag{S104}$$

where $n_i$ is the number of cells of strain $i$, and $\lambda_i$ is the corresponding growth rate is not enough. Instead, we will assume that the cells follow the logistic growth equation of the form

$$\frac{dn_i}{dt} = \lambda_i n_i \left(1 - \frac{\sum_{j=1}^{N} n_i}{\kappa}\right), \tag{S105}$$

where $\kappa$ is the carrying capacity, and $N$ is the total number of strains in the culture.

The inference method is based on the model that assumes that the time passed between dilutions $\tau \approx 8$ generations, the change in frequency for a mutant barcode can be approximated from cycle $t$ to the next cycle $t+1$ as

$$f_{t+1}^{(m)} = f_t^{(m)} \mathrm{e}^{(s^{(m)} - \bar{s}_t)\tau}, \tag{S106}$$

where $s^{(m)}$ is the relative fitness for strain $i$ compared to the ancestral strain and $\bar{s}_t$ is the mean fitness of the population at cycle $t$. To test this assumption, we implemented a numerical experiment following the logistic growth model described in Equation S105. Figure S2 shows an example of the deterministic trajectories for 50 labeled neutral lineages and 1000 lineages of interest. The upper red curve that dominates the culture represents the unlabeled ancestral strain included in the experimental design described in Section .

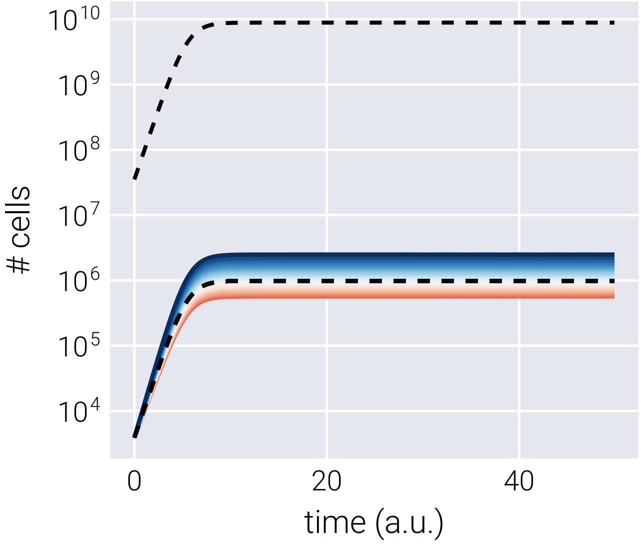

**Figure S2. Logistic growth simulation over single growth cycle**. The dashed line represents the neutral lineages, with the upper curve being the unlabeled neutral strain. Color curves represent the genotypes of interest colored by growth rate relative to the neutral lineage.

To simulate multiple growth-dilution cycles, we take the population composition at the final time point and use it to initialize a new logistic growth simulation. Figure S3 shows the resulting number of cells at the last time point of a cycle over multiple growth-dilution cycles for the genotypes in @Figure S2. We can see that the adaptive lineages (blue curves) increase in abundance, while detrimental lineages (red curves) decrease.

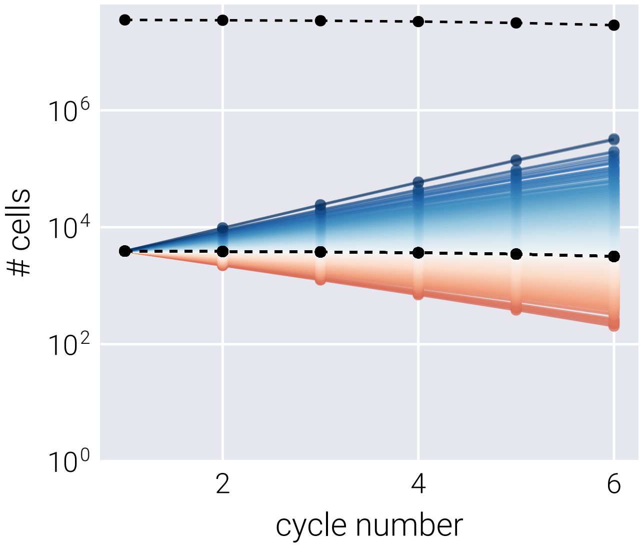

**Figure S3. Growth-dilution cycles for logistic growth simulation**. Each point represents the final number of cells after a growth cycle for each lineage. Colors are the same as in Figure S2.

In Section , we derive the functional form to infer the relative fitness of each lineage as

$$\frac{1}{\tau} \ln \frac{f_{t+1}^{(b)}}{f_t^{(b)}} = (s^{(b)} - \bar{s}_t). \tag{S107}$$

Figure S4 shows the corresponding log frequency ratio curves for the logistic growth simulation. The displacement of these curves with respect to the neutral lineages determines the ground truth relative fitness value for these simulations.

To simulate the experimental noise, we add two types of noise:

1. Poisson noise between dilutions. For this, we take the final point of the logistic growth simulation and sample a random Poisson number based on this last point to set the initial condition for the next cycle.
2. Gaussian noise when performing the measurements. When translating the underlying population composition to the number of reads, we can add a custom amount of Gaussian noise.

Figure S5 shows the frequency trajectories (left panels) and log frequency ratios (right panels) for a noiseless simulation (upper panels) and a simulation with added noise (lower panels). The noiseless simulation is used to determine the relative fitness for each of the lineages, which serves as the ground truth to be compared with the resulting inference.

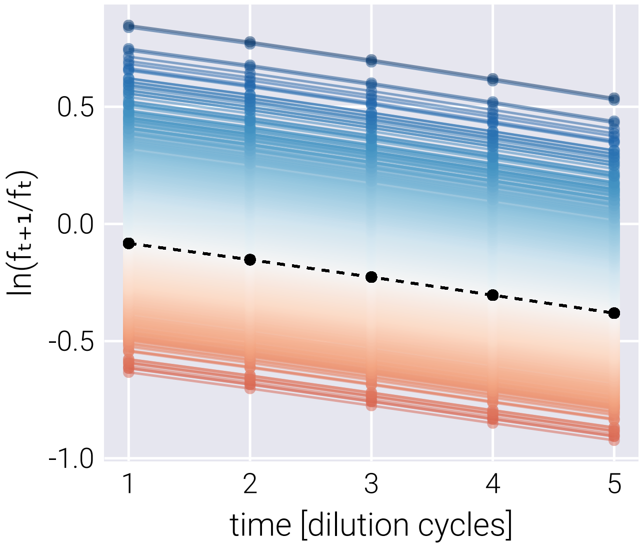

**Figure S4. Log frequency ratio for logistic growth simulations**. The relative distance of the color curves from the black dashed line determines the relative fitness of each linage.

1021 # Supplemental References

1022 [25]A. Gelman, D. Simpson, and M. Betancourt, "The Prior Can Often Only Be Understood in
1023 the Context of the Likelihood", Entropy **19**, 555 (2017).

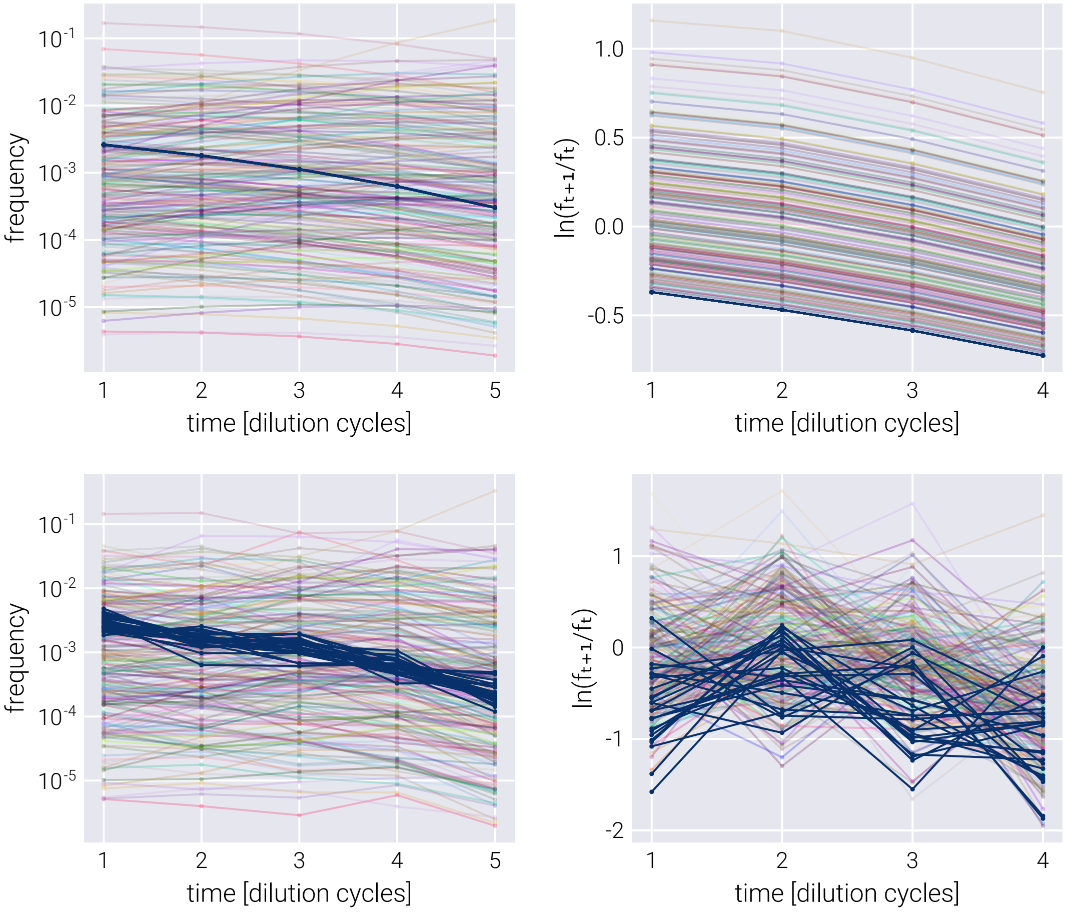

**Figure S5. Logistic growth-dilution simulations with and without noise**
